# Illumination optimization and low-power trapping of *Limnospira indica* PCC 8005 using bulk acoustic waves in microgravity

**DOI:** 10.1101/2025.05.27.656267

**Authors:** Bérénice Dupont, Xavier Benoit-Gonin, Sébastien Vincent-Bonnieu, Jean-Luc Aider, Maxime Ardré

## Abstract

Space missions require sustainable life support systems capable of producing oxygen and biomass under microgravity. We report the use of acoustic levitation to trap and manipulate the filamentous cyanobacterium *Limnospira indica* PCC 8005 during parabolic flights. Within a millimeter-scale fluidic chamber, this helical microorganism rapidly assembles into thin layers under a standing ultrasonic wave. Stable trapping in microgravity requires substantially less acoustic power (0.42 mW) than on Earth (1.4 mW), highlighting the potential for energy-efficient bioprocessing in space. Monte Carlo simulations and light attenuation modelling show that layered structuring enhances light penetration, potentially overcoming the “compensation point” limitation in bulk cultures. These findings open new perspectives for photobioreactors using acoustic manipulation to boost photosynthetic efficiency and reduce energy demands for oxygen and biomass production in space.

## Introduction

Space exploration and human spaceflight have long been central goals for national space agencies, but their importance has increased over the past decade with the emergence of non-governmental companies such as SpaceX and Blue Origin. A key objective is to enable long-term space missions and establish permanent extraterrestrial bases, first on the Moon (NASA’s Artemis program) and later on Mars (SpaceX), both of which depend on the development of advanced environmental control and life support systems^1^. Over the past 30 years, significant progress has been made in recycling carbon dioxide (CO_2_) and water aboard the International Space Station (ISS)^2,3^. In addition, the European Space Agency (ESA) launched the MELiSSA (Micro-Ecological Life Support System Alternative) project in 1987 to design and develop a bioregenerative closed-loop life support system for long-duration missions, aimed at supplying food, oxygen, potable water and hygienic water while efficiently recycling waste^4–7^.

Microalgae and cyanobacteria are promising candidates for such systems due to their photosynthetic capabilities^8^. Cyanobacteria are photosynthetic prokaryotes capable of converting CO_2_ into biomass while simultaneously releasing O_2_^9^. Among them, *Limnospira indica* PCC 8005 is a well-established model organism in the MELiSSA project^10,11^. In liquid culture, this cyanobacterium forms multicellular filaments, or trichomes, composed of cells joined end-to-end. Its adaptability to space conditions is well documented^12,13^. It can serve a dual role: (i) as a renewable biological system for O_2_ production via photosynthesis and (ii) as a source of edible biomass^14^, provided that light and carbon availability are sufficient to sustain growth. Optimizing photobioreactors is therefore essential for efficient cultivation.

A photobioreactor is a closed system designed to grow light-dependent organisms under controlled conditions. In such systems, light absorption and scattering determine how light propagates through the culture and reaches every microorganism. Light absorption by the culture causes an exponential decrease of light intensity as a function of the light-path in the fluid (Beer–Lambert law), while scattering redirects light in different directions when it encounters particles or microorganisms. Several models have been developed to describe light penetration in cultures of photosynthetic microorganisms, accounting for both absorption and scattering^15–17^. These models show that incident light is typically extinguished within the first few centimeters of the culture.

The so call *compensation point* quantifies this limitation^16^. It is defined as the distance from the light source where light intensity drops below the minimum required for photosynthesis. Beyond this point, photosynthetic activity ceases, limiting biomass production and CO_2_ conversion to O_2_. The position of the compensation point depends on cell concentration, photobioreactor geometry, illumination configuration, and the optical properties of the cells. Thus, increasing photosynthetic yield requires strategies to shift the compensation point deeper into the culture, exposing more cells to light.

Acoustic levitation provides contact-free control of microscopic objects in fluids. A standing ultrasonic wave, generated by a transducer in a resonant cavity, produces an acoustic radiation force (ARF) that drives suspended particles toward the pressure nodes of the field^18,19^. Since the node pattern is fixed, the ARF precisely localizes particles along the cavity height. Previous studies have demonstrated ARF-based trapping of spherical objects such as micrometric particles, cells, and microalgae^20–24^. However, controlling the spatial distribution of living cyanobacteria for functional purposes such as improving light access remains unexplored. Trapping efficiency depends on factors such as particle size, transducer voltage and frequency, and external forces (gravity, fluid acceleration). Most ARF models apply to inert, spherical^18,20,25–28^ or elongated^29–33^ particles small compared to the acoustic wavelength (Rayleigh regime). By contrast, *L. indica* PCC 8005 trichomes are thin, helicoidal, soft, biologically active and relatively long compared to the acoustic wavelength (about 740 µm at 2.00 MHz in water) with lengths ranging from a few hundred micrometers to several millimeters^9,34^) thus deviating substantially from model assumptions. Although chemically fixed *L. indica* has been levitated^34^, the response of live cells remains uncertain, as ongoing metabolism alters cytoplasmic composition, density, and compressibility. For space applications, two key questions arise: (i) Can ARF effectively trap living *L. indica* PCC 8005? and (ii) What electrical power is required to achieve stable trapping under varying gravity conditions?

Here we show that a standing ultrasonic wave can gather *L. indica* at pressure nodes under these experimental conditions, stacking them into thin, cell-dense layers separated by clear fluid gaps. These gaps act as optical windows, enabling light to penetrate to depths that remain dark in fully mixed suspensions. Monte Carlo photon-transport simulations confirm that this layered arrangement increases fluence rates throughout the reactor, enhancing light availability to more cells. We quantify trapping performance in a flight-qualified millifluidic chip, measuring the drive voltage required to form and maintain the layers at 0 g during parabolic flights and at 1 g in the laboratory. In microgravity, the electrical power threshold is roughly half that on Earth, indicating that milliwatt-level input is sufficient for continuous operation in space. These results identify acoustic levitation as a low-energy method for structuring cyanobacterial cultures, advancing their potential for future life-support systems.

## Methods

### Acoustic Levitation in Resonant Cavities

An acoustic standing wave (ASW) within a resonant cavity makes possible the trapping of objects in a liquid medium without any physical contact. Figure 1 illustrates the core principles of this technique. If an ultrasonic transducer located at the top of the cavity emits an acoustic wave inside the cavity toward the lower glass wall, which behaves as a rigid reflector, and if the acoustic wavelength *λ* is chosen, such as the cavity height *h* is a multiple of half wavelength 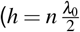, where *n* ∈ ℕ^∗^), then the resonant condition is achieved^18^ and an ASW is formed (Fig. 1b). In the following, the resonant wavelength and corresponding frequency are noted *λ*_0_ and *f*_0_. When this condition is satisfied, one (*n* = 1) or multiple (*n >* 1) pressure nodes appear along the cavity’s height, spaced at intervals of Δ*z* = *λ*_0_*/*2, where *z* is the axial direction corresponds to the direction of propagation of the wave, perpendicular to upper and lower walls.

**Figure 1:**
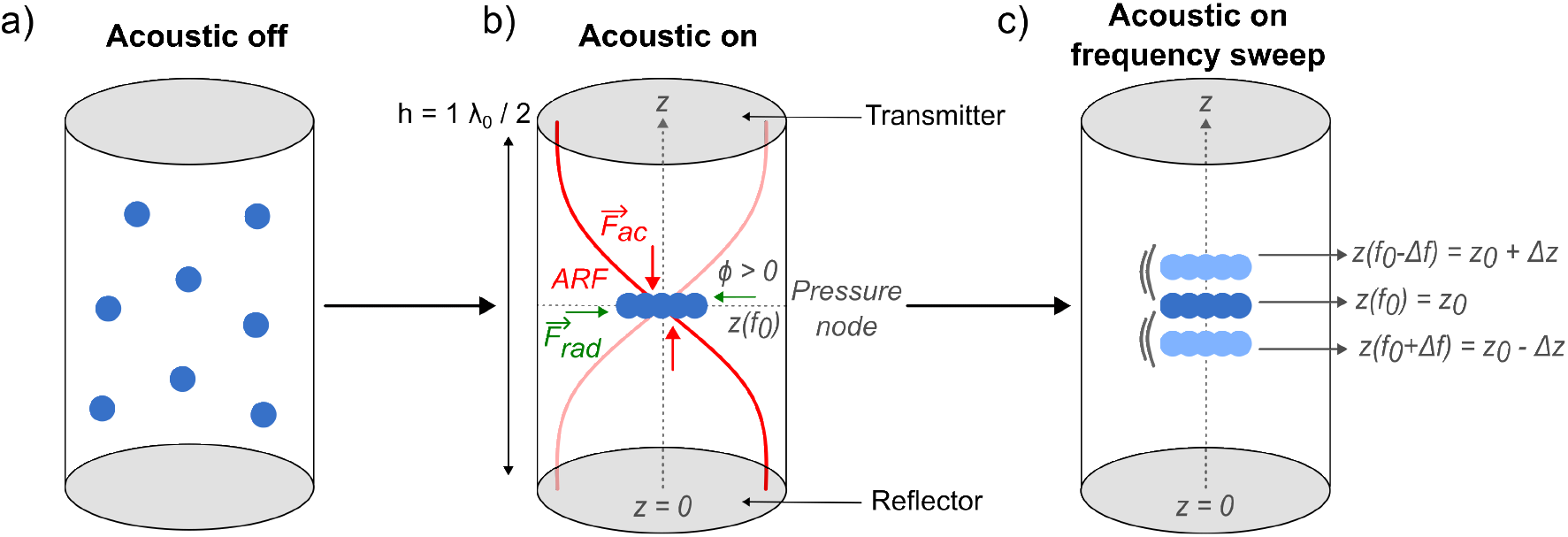
Principle of Acoustic Levitation in a Resonant Cavity. **a)** Particles (blue) with a positive contrast factor (*ϕ >* 0) are initially uniformly distributed in the fluid-filled cavity. **b)** When the ultrasonic transducer is activated at the resonance frequency *f*_0_, the particles migrate toward the pressure node and aggregate at this stable levitation plane. In this example, *h* = *λ*_0_*/*2. **c)** Varying the acoustic frequency near resonance (*f*_0_) leads to an axial displacement of the levitation plane. A frequency change Δ *f* corresponds to a displacement Δ*z* of the levitation plane, enabling precise vertical positioning of the trapped particles. A frequency sweep leads to an axial oscillation of the objects trapped in the levitation plane.

Let’s consider small spherical objects initially homogeneously distributed in the volume of the fluidic cavity (Fig. 1a). Once the ASW is established, any object with physical properties, like density or compressibility, sufficiently different from the suspending fluid experiences the so-called Acoustic Radiation Force (ARF). For a spherical, compressible particle much smaller than the acoustic wavelength, i.e. *d*_*p*_ ≪ *λ* (with *d*_*p*_ the diameter of the object), the axial component of ARF *F*_ac_ can be expressed as^25,35^:

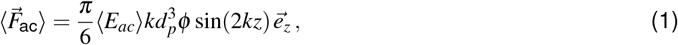

where ⟨−⟩ represents time average, *E*_*ac*_ is the acoustic energy density, *k* = 2*π/λ*_0_ is the wavenumber, *ϕ* is the acoustic contrast factor, and *z* is the axial position of the particle.

The acoustic contrast factor (ACF) *ϕ* characterizes the difference in mechanical properties between the particle and the surrounding fluid:

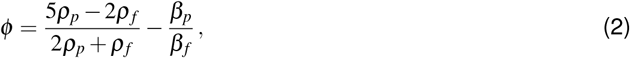

where *ρ*_*p*_ and *ρ*_*f*_ are the mass densities of the particle and fluid, respectively, and *β*_*p*_ and *β*_*f*_ are their respective compressibility. Particles with a positive ACF (*ϕ >* 0) are driven towards the closest acoustic pressure node of the ASW (as illustrated with the force 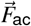 in Fig. 1b. Once they have reached the closest pressure node, they can be maintained in acoustic levitation as long as needed. It should be noted that objects with a negative ACF, like oil droplets, will move toward the acoustic pressure anti-nodes. Typical forces range from a few pN to about 0.01 pN^36^. For instance, a 10 µm diameter polystyrene bead in water (*ρ*_*p*_ = 1.05 g.cm^−3^, *β* = 1.6 × 10^−10^ Pa^−1^, *ϕ* = 0.24) subjected to an acoustic energy density of 10 J.m^−3^ experiences an ARF of about 2×10^−12^ N (2 pN). This force is one order of magnitude larger than the apparent weight of the bead. Indeed, the buoyancy force is A = 5.13 pN and the weight is W = 5.39 pN, leading to an apparent weight W-A = 0.79 pN.

In a second step, once the particles have reached the levitation plane, they are slowly drawn toward the centre of the cavity by the radial component of the ARF, noted 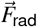. Because this component is much weaker (10 to 100 times weaker) than the axial component of the ARF^37^ it is often neglected. Nevertheless, it is strong enough to move the particles inside the levitation plane toward the centre of the cavity where they can form aggregates.

Two key acoustic parameters govern the effectiveness of acoustic levitation: first, the acoustic energy density, ⟨*E*_*ac*_⟩ which depends on the square of the voltage *U*_*ac*_ applied to the transducer 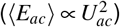 up to the transducer’s operational limits^38,39^, second, the acoustic wavelength *λ* (or equivalently the acoustic frequency *f* × *λ* = *c, c* being the speed of sound in the fluid medium), which determines the axial position of the pressure nodes if the resonance condition is satisfied.

Within a multi-layered acoustofluidic chip (a transmitting layer on the top and a reflecting layer on the bottom of the cavity), studies have demonstrated^35,40–43^ that varying the acoustic frequency by a few tens of kHz around the 2.0 MHz resonance frequency, the *z*-positions of the levitation planes can be shifted along the longitudinal (*z*) axis, either with a single node^35^ or multinode cavity^43^. In our device we show that the *z*-positions is proportional to the frequency of the transducer Fig. S9. In the same time, the distance between two levitation planes is also modified (reduced when the frequency is increased). It should be noted that these experiments require a wide-band acoustic transducer so that the acoustic energy emitted by the transducer decreases slowly around its resonance frequency. Using this property, small variations of frequency Δ *f* around the resonance frequency *f*_0_ lead to small variations Δ*z* around the node axial position at the resonance *z*_0_, as shown in Fig. 1c and Fig. S9. By sweeping the frequency within a range [*f*_0_ −Δ *f, f*_0_ +Δ *f*], one can move objects in levitation along the z-axis around the equilibrium position at the resonance ([*z*_0_ + Δ*z, z*_0_ −Δ*z*]). This oscillation of the levitation plane is possible only if the axial ARF *F*_*ac*_ is strong enough over the chosen frequency range.

By adjusting both the voltage applied to the transducer (*U*_*ac*_) while varying the acoustic frequency *f* near *f*_0_ it becomes possible to identify the minimum electrical power needed to reliably trap cyanobacteria in acoustic levitation.

### Millifluidic chip

The chip is made of a molded PDMS (polydimethylsiloxane, PDMS Sylgard 184) bonded to a microscope glass slide (reflecting layer). The design of the milli-fluidic chip is shown in Fig. 2a. The inner dimensions of the cavity are 14 mm × 10 mm × 3 mm (length × width × height). The key feature of the chip is the height of its central cavity h_1_ (Fig. 2b). The total height (H) of the chip corresponds to the sum of the cavity height h_1_ and the height of the PDMS layer h_2_ (transmitting layer) of about 0.7 mm, leading to H = 3.7 mm. The thickness of the microscope slide (reflecting layer) is 1 mm. The coupling between the PDMS chip and the transducer is achieved by a thin layer of oil, as illustrated in Fig. 2b. The fabrication of the millifluidic chip begins with the preparation of a 10:1 mixture of PDMS elastomer base and curing agent. This mixture is subjected to vacuum degassing to remove any trapped air bubbles. After degassing, the mixture is carefully poured into moulds and cured at 70°C for 2 hours. Once curing is complete, the chips are gently extracted from the moulds. Inlet and outlet wells are manually punched using a 2.0 mm diameter piece cutter. Both the PDMS chips and microscope glass slides are cleaned with isopropanol to ensure optimal adhesion. Finally, the PDMS pads are bonded to the glass slides via plasma treatment (20 W, 50 Hz, 8 sscm O_2_).

**Figure 2:**
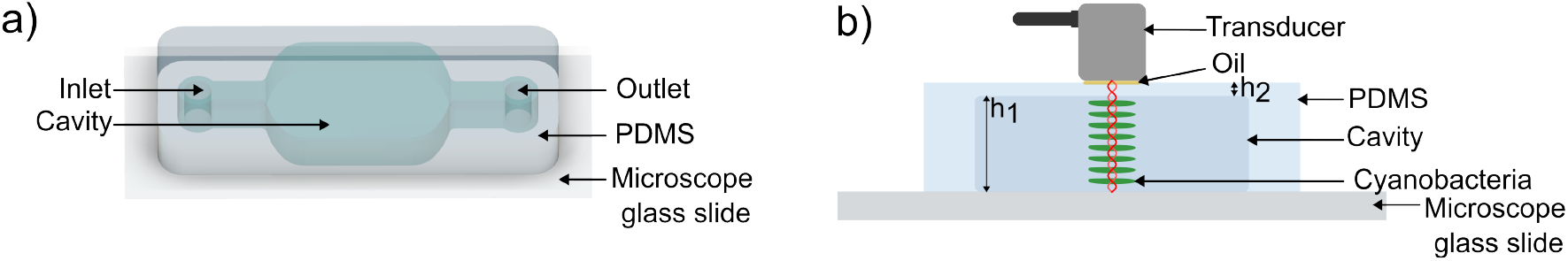
Acoustofluidic chip design. **a)** CAD model of the chip, showing the inlet, outlet, and central cavity. **b)** Schematic of the millifluidic chip containing a cyanobacterial culture organized in layers localized at the nodes of the resonant cavity. The transducer is mounted above the cavity. The red waving lines depict the acoustic standing waves in the cavity. The green layers illustrate the aggregation of the cyanobacteria that cluster at the nodes of the wave. A typical scale is given by h_1_ which is 3 mm long.

Cyanobacteria suspended in a nutrient medium are introduced through the inlet into the central cavity of the chip using a pipette tip. The inlet and outlet are then sealed with PDMS pads.

### Acousto-fluidic setup

A compact Signal Processing™ (TR0205RSX) 2.0 MHz acoustic transducer, with a diameter of 5 mm and a body length of 21 mm, is placed on the top of the cavity to generate the ultrasonic wave. A thin layer of oil between the transducer and the PDMS is used to match their impedance. The transducer is powered by a Tie-Pie USB wave generator (Handyscope HS5 XM). Both the voltage and the frequency are controlled using the Tie-Pie software. The electrical power consumption is monitored by the TiePie software during the operations. For a 2.0 MHz acoustic frequency, the acoustic wavelength *λ*_0_ is about 740 µm in water at 20°C, which leads to the formation of 8 pressure nodes along the cavity’s height (*h*_1_). Once the ASW is formed, the objects in suspension migrate toward the closest pressure node, as illustrated in Fig. 2b, and form layers.

To evaluate the efficiency of the acoustic trap as a function of the acoustic energy density (proportional to 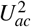), a linear frequency sweep is applied from 1.8 to 2.2 MHz with a 450 ms duration cycle. The frequency sweep leads to the oscillation of the axial position of the clusters, provided that the acoustic energy density is high enough.

### Experimental setup in the AirZeroG plane

The setup, adapted from previous studies^32,44^, is securely installed in a waterproof Zarges box that is firmly mounted inside the aircraft (Fig. 3a). To prevent fluid leakage, the setup is kept inside the box during the flights. A 3D-printed structure has been designed to securely hold the transducer, the chip, and a Dino-Lite™ mini-microscope (AM2111) for side observations of the chip (Fig. 3b). The wave generator and the digital microscope are controlled by a laptop mounted on a dedicated rack (Fig. 3a). The other elements shown in Fig. 3a are used for another unrelated experiment^44^, which is not discussed in this article.

**Figure 3:**
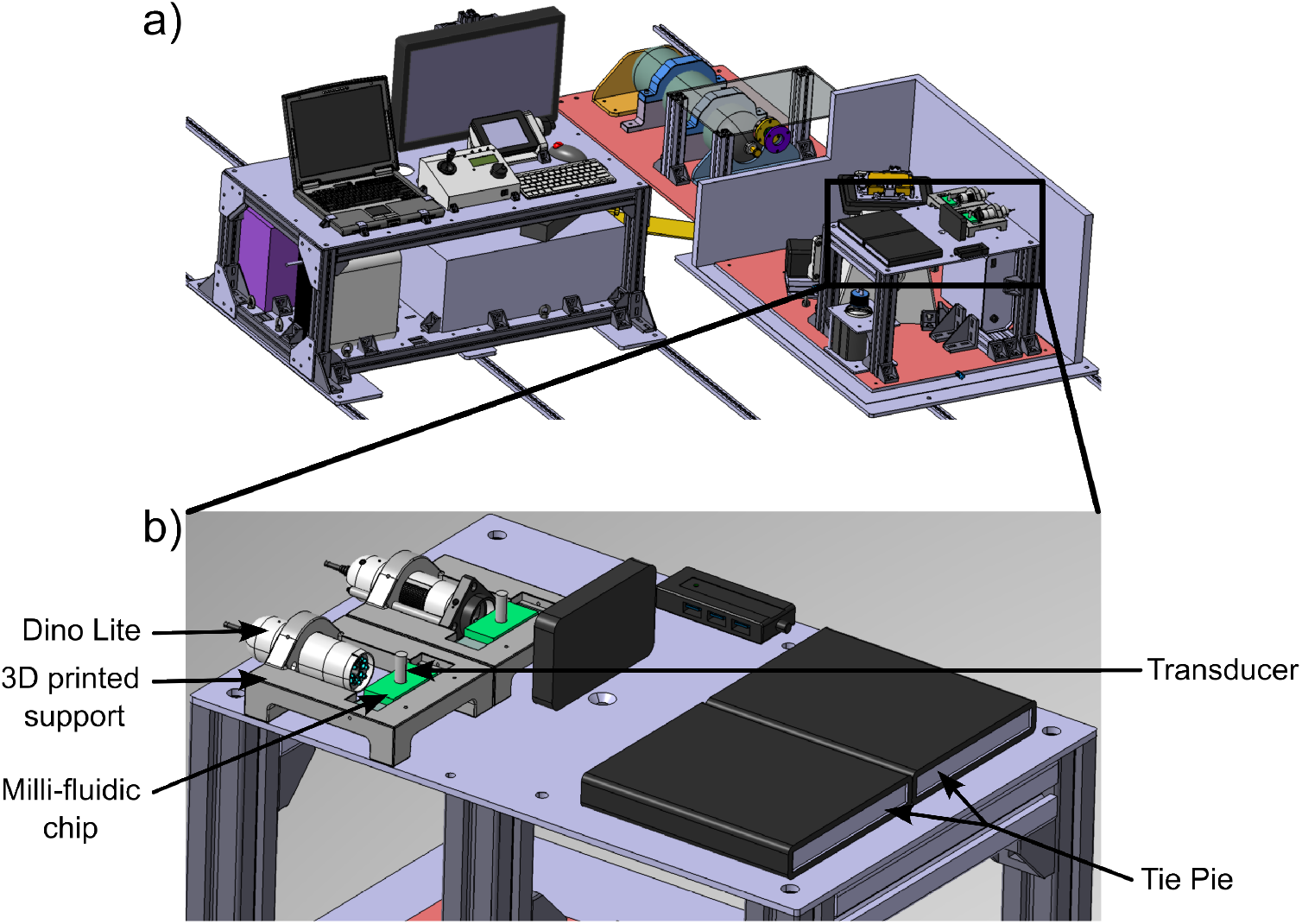
3D view of the experimental setup. **a)** Full view of the experimental setup. It includes a control rack (on the left), which allows the control of all the experimental parameters. A sealed Zarges box contain the experimental setup (on the right). Two experimental devices were installed in the box: a Zeiss inverted microscope (on the left side) for cell biology experiments^44^ and the milli-fluidic chips to study the acoustic trapping of cyanobacteria (on the right side). **b)** Zoom on the two PDMS chips that were used to trap the cyanobacteria. Digital microscopes (Dino Lite) were used to monitor the aggregation during the parabolas.

Each parabolic flight typically consists of six sets of five parabolas, for a total of 30 periods of micro-gravity of about 22 s each. Throughout the flight, the box can be opened a few minutes after take-off before the first parabola for about 5 min and then six times at every break of 5 min in between the series of parabolas. This constraint prevents any modification to be applied to the device during the experiment (for example, shaking or refilling).

### *Limnospira indica* PCC 8005

The multicellular, filamentous cyanobacteria *Limnospira indica* PCC 8005 was purchased from the Pasteur Culture Collection (PCC). The organisms were cultivated in 150 ml of a nutritive Zarrouk’s medium (Tab. 1) under aerobic conditions at 27°C, with an illumination intensity of 6.64 W · m^−2^ (equivalent to 2500 lux) measured at 20 cm from the light source. Flasks were incubated in a rotary shaker set at 140 rpm. The experiments were conducted with an optical density (OD) of approximately 0.1, measured at 750 nm (optical wavelength). When needed, a sample is taken from a batch culture and injected into the chip to perform the experiments.

**Table 1:** Zarrouk medium^45^ composition, including trace elements.

| Composition of the culture medium | Concentration (g·L<sup>-1</sup>) |
| --- | --- |
| NaCl | 1.0 |
| CaCl<sub>2</sub> | 0.04 |
| K<sub>2</sub>HPO<sub>4</sub> | 0.5 |
| NaNO<sub>3</sub> | 2.5 |
| NaHCO<sub>3</sub> | 5.0 |
| K<sub>2</sub>SO<sub>4</sub> | 1.0 |
| MgSO<sub>4</sub> | 0.1 |
| FeSO<sub>4</sub> | 0.01 |
| EDTA | 0.08 |
| Traces elements (A5) with Co | 1 mL |
| Traces elements (A5) with Co | Concentration (g·L<sup>-1</sup>) |
| H<sub>3</sub>BO<sub>3</sub> | 2.86 |
| MnCl<sub>2</sub> | 1.81 |
| ZnSO<sub>4</sub> | 0.222 |
| NaMoO<sub>4</sub> | 0.390 |
| CuSO<sub>4</sub> | 0.079 |
| Co(NO<sub>3</sub>)<sub>2</sub> | 0.049 |

In this work OD_750_ is used as a convenient proxy to measure biomass as it is linearly proportional to the trichome biomass, directly measured with a microscope, as shown in Fig. S8.

The morphological characterization shown in Fig. 4b was performed on *Limnospira indica* PCC 8005 filaments (or trichomes) obtained from three independent cultures, pooled together, with an optical density 0.2 (OD_750_). For each culture, 25 filaments were imaged Fig. 4a and measured, totalling 75 filaments. In each case, measurements were made on three separate samples per culture. Observations were carried out using a Zeiss Imager Z2 microscope at 5X in bright-field with a Mallassez counting chamber. Filament (or trichomes) lengths and widths were measured using ImageJ (Fiji). Under these experimental conditions, the cyanobacterial filaments of *L. indica* PCC 8005 exhibits a spiral, rod-like morphology (Fig. 4a), with lengths ranging from 100 µm to 200 µm and a mean width of 25 µm shown Fig. 4b.

**Figure 4:**
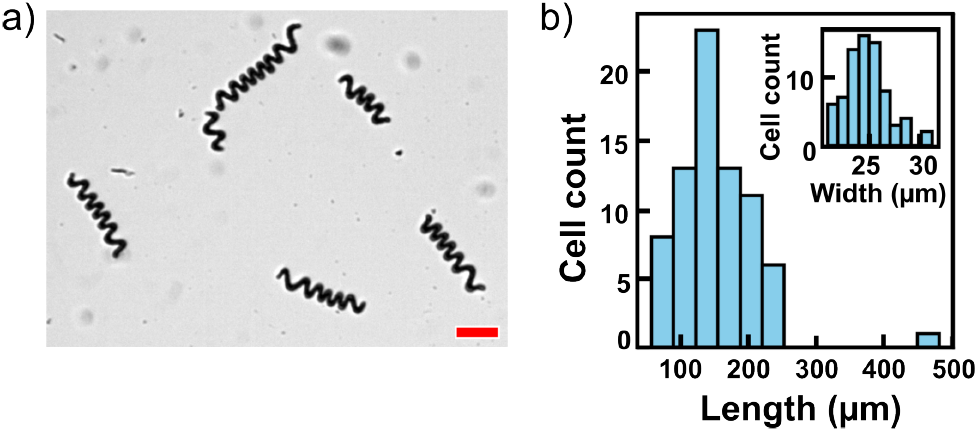
Characterization of *L. indica*. **a)** Micrograph of *L. indica* PCC 8005 filaments. Scale bar, 50 µm. **b)** Size distribution (length and width) of *Limnospira indica* PCC 8005 in liquid suspension (n = 75).

The acoustic contrast factor of the cyanobacteria was assessed at 2.00 MHz and 10 V in the condition of the 0 g experiments. They were mixed with 300 µm diameter polystyrene beads with positive acoustic contrast factor (that localized at the nodes). The experiment was carried out in the chip shown in Fig. 2, which has an internal height *h*_1_ of 4 mm (unlike the ones used during the flights for which *h*_1_ is 3 mm).

### Image analysis

The displacements of the levitated layers were recorded with a Dino-Lite AM2111 USB microscope (640 × 480 px, 20 fps) rigidly fixed to the millifluidic chip holder. AVI stacks were imported into Fiji as 8-bit grayscale images, with the *z*-coordinate defined in ImageJ by drawing a line perpendicular to the transducer surface. The images were then cropped to a ∼45-px-wide window centered on the targeted layer. After applying a binary threshold, a 20-pixel-wide region of interest (ROI) was defined along the axial direction (*z*-axis) through the layer’s midpoint to extract an axial intensity profile. For the analysis, only one aggregate was selected, since most microgravity data did not consistently allow tracking of multiple aggregates due to successive 0 g-2 g phases without manual intervention. The *z*-coordinate of the uppermost dark pixel within this line representing the highest point of the cyanobacterial layer was measured for each frame, yielding the time evolution of the axial position *z*(*t*) of the aggregate. Figure S2 illustrates the processing steps.

When the ARF was sufficiently strong, *z*(*t*) exhibited periodic oscillations. For each driving voltage *U*_ac_, the mean peak-to-peak amplitude was extracted, and the standard deviation computed over 11 s of recording. Error bars in Fig. S3 represent the standard deviation from more than 20 measurements per voltage value. This repetition enabled estimation of the mean oscillation amplitude with a precision exceeding the 12 µm pixel resolution, since the distribution of values allows sub-pixel accuracy under the assumption of Gaussian noise. Nonetheless, the limit of detection of 12 µm was used as the threshold to classify “no oscillation”. Thus, the mean amplitudes below 12 µm were set to 0 µm.

### Electrical power measurements

The electrical power delivered to the piezoelectric transducer was measured using the Tie-Pie USB Handyscope and its Multi-Channel v1.45.1 software. Power consumption was measured directly using the Power function from Meter function within the software, which provides an estimation of the active electrical power delivered to the transducer during operation. Measurements were carried out on two different PDMS acoustofluidic chips: the millifluidic chip described above, with an internal volume of cm^3^ (3 mm-high), and a chip twenty times larger, with an internal volume of 8 cm^3^ (8 cm-high × 1 cm-wide), previously designed in the study of Bellebon *et al*.^24^.

## Results

### Acoustic levitation of cyanobacterial trichomes

The acoustic trapping of micrometric particles is well described by the theory, which assumes that particles are spherical and much smaller than the acoustic wavelength (740 µm in the fluid medium and *f* = 2 MHz)^21,35,46^.

The effective trapping of living *Limnospira indica* PCC 8005 by the ARF is demonstrated in Fig. 5. The cyanobacterial culture was injected into the millifluidic chip (Fig. 2) before activating the ultrasonic transducer. Fig. 5a–c shows that the clustering process occurs in two distinct steps. Initially, the strongest *z*-component of the ARF (*F*_ac_) acts on the suspended cyanobacteria (Fig. 5b), rapidly forming (within s) multiple layers at the pressure nodes of the multinode resonant cavity. Subsequently, after approximately 15 s (Fig. 5c), aggregates begin to form at the center of each layer, driven by the weaker radial component of the ARF (*F*_rad_). This results in compact, layer-like clusters with a thickness along the *z*-axis of about 100 µm and a lateral width of approximately 1 mm. These observations demonstrate that living, helicoidal cyanobacteria can be stably trapped, layered, and repositioned with acoustic forces, despite their complex geometry and time-varying material properties.

**Figure 5:**
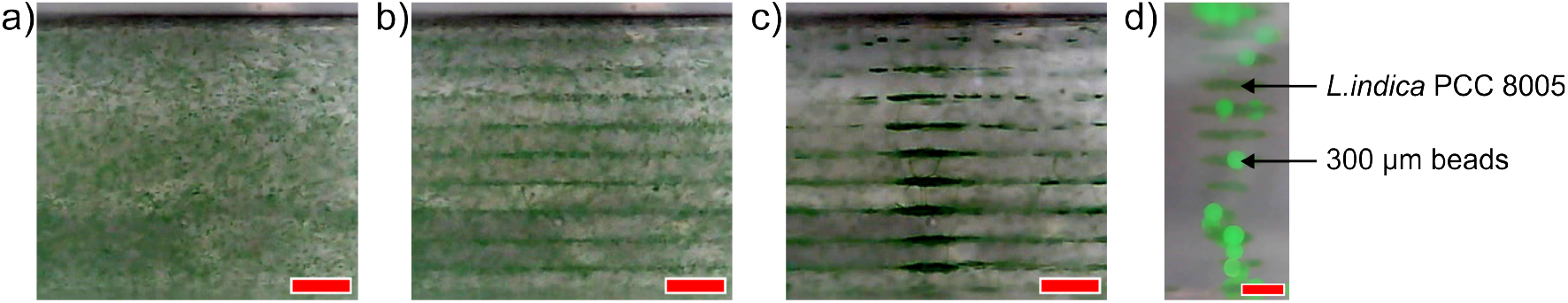
Side observation showing the formation of layers of *Limnospira indica* PCC 8005 using acoustic levitation in terrestrial condition. **a)** Cyanobacteria suspension prior to acoustic field activation. **b)** Formation of layers of cyanobacteria 0.5 s after activation of the acoustic field. **c)** Formation of clusters of cyanobacteria in the central region of the cavity 15 s after activation (Movie online provided in supplementary data). Scale bars, 500 *µ*m. **d)** *L. indica* PCC 8005 mixed with 300 µm diameter green polystyrene beads. One can see that the cyanobacteria form clusters on the same axial position than the beads. The scale bar is 500 *µ*m.

Cyanobacteria such as *L. indica* possess gas vesicles, so their mass density may vary over their life cycle. In our conditions, before the parabolic flights, the sign of the acoustic contrast factor (ACF) of *L. indica* PCC 8005 was evaluated by mixing a suspension of cyanobacterial filaments with 300 µm polystyrene beads in a millifluidic device (with *h*_1_ = 4 mm). The polystyrene beads have a positive ACF and thus serve as a reference for node localization. Fig. 5d shows that both the cyanobacterial filaments and the polystyrene beads localize at the same horizontal planes corresponding to the pressure nodes of the standing wave. Thus, under these conditions, *L. indica* exhibits a positive ACF. Besides, the spacing between the levitation planes is close to *λ*_0_*/*2 ≈ 370 *µ*m, consistent with the bead diameter.

Furthermore, although previous studies reported destructive effects on cyanobacteria^47^ in sonication setups producing progressive pressure waves of about 1 m wavelength, this is not the case here for a stationary pressure wave in a resonant microcavity with an acoustic wavelength of 740 µm. Fig. S1 (supplementary data) shows that cellular viability is not affected by acoustic levitation. A culture levitated for 6 h displayed growth comparable to a standard culture, and the morphology of the cyanobacteria remained unchanged after 1 h of acoustic levitation.

Therefore, the spontaneous formation of alternating layers of biomass and transparent medium is possible, even though the system deviates significantly from ideal theoretical conditions. This self-organization offers a promising strategy for improving cyanobacterial illumination in photobioreactors.

### Acoustic layering to enhance light penetration in cyanobacterial photobioreactors

Conventional photobioreactors rely on suspended cultures that are regularly agitated to expose all cells to light. Here, we propose using acoustic radiation forces (ARF) to arrange the cyanobacterial culture into layers to increase light penetration into the reactor.

To evaluate whether such acoustic layering could mitigate the *compensation point* limitation^16^, photon-transport simulations were carried out using *MCmatlab*^48^. Two geometries were compared:

1. Bulk: the cells are uniformly distributed at *C*_*x*_ = 1 kg·m^−3^;
2. Leaf: the cells are organized into three layers, each 100 µm thick at *C*_*x*_ = 3.33 kg·m^−3^, separated by transparent fluid (Fig. 6b), mimicking the configuration observed experimentally (Fig. 5).

**Figure 6:**
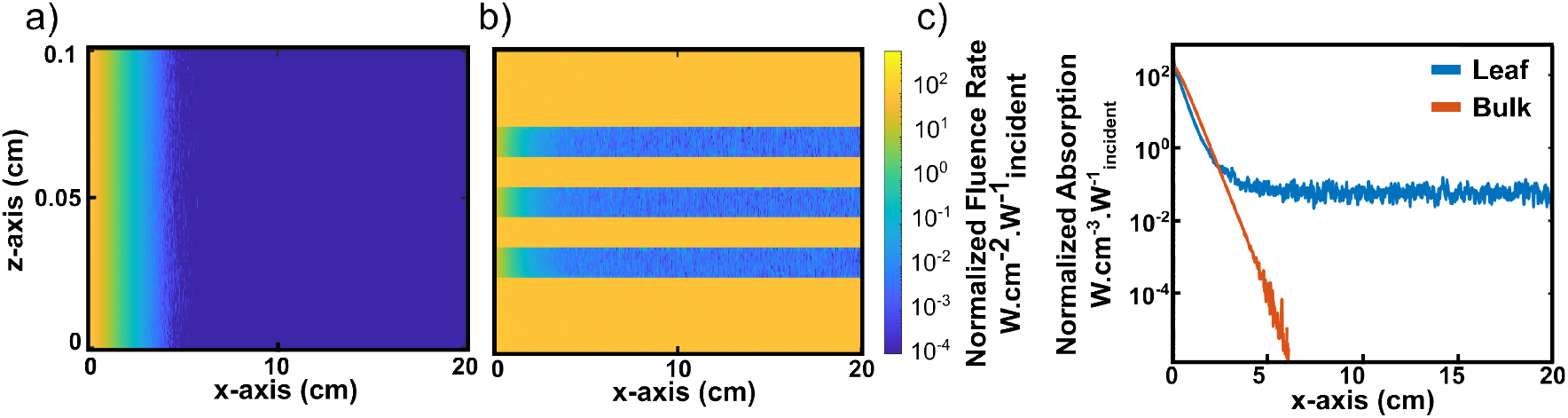
Monte Carlo simulations of light distribution in a photobioreactor with and without layering of cyanobacterial culture. **a)** Fluence rate of light in a vertical slice of a bulk culture where cyanobacteria are suspended. The light source is located at *x* = 0. **b)** Fluence rate of light in a vertical slice when cyanobacteria are trapped, forming three distinct layers interspaced by transparent fluid. The z-axis and x-axis are scaled differently for readability; a) and b) share the same z-axis. **c)** Mean normalized absorption profiles along the x-axis for homogeneously distributed (bulk) and layered (leaf) cultures.

In the Leaf case, the local cell concentration within the layers is higher than in the Bulk configuration because the same total biomass is confined within a smaller volume (0.3 times smaller). Both simulations used identical optical parameters (*E*_*a*_, *E*_*s*_, and *g*) from the Cornet model^49^. A top-hat beam enters at *x* = 0 with a slight angle (arctan(*L*_*z*_*/L*_*x*_)) and propagates across the domain (*L*_*z*_ = *L*_*y*_ = 0.1 cm, *L*_*x*_ = 20 cm). Light exiting at *x* = *L*_*x*_ is considered lost. Cyclic boundary conditions on the planes *z* = 0, *z* = *L*_*z*_, *y* = 0, and *y* = *L*_*y*_ allow photons exiting one side to re-enter from the opposite side.

Figures 6a,b show the resulting fluence-rate fields normalized by the incident flux. Figure 6c shows the mean absorbed power density normalized by the incident flux as a function of the light path. This quantity is averaged over the full cavity height in the Bulk case, and only within the layers in the Leaf case.

In the Bulk configuration, light flux drops to near zero within the first 6 cm (Fig. 6a), consistent with the classical *compensation point* bottleneck. In contrast, in the Leaf configuration (Fig. 6b), the light flux decays more slowly, stabilizing beyond *x* = 4 cm because photons crossing the transparent gaps can still excite downstream layers. Thus, layering is expected to improve the overall light distribution deeper inside the photobioreactor and potentially keeps a larger fraction of cyanobacteria above the photosynthetic threshold. Moreover, despite their higher biomass concentration, the layers still permit light penetration, as evidenced by their uniform light-blue coloration in Fig. 6b, which corresponds to a significant fluence rate. This indicates that self-shading within each layer is minimal.

These simulations demonstrate that acoustic layering can achieve a more uniform light distribution deeper in the photobioreactor, which is a critical factor for maximizing photosynthetic efficiency in energy-limited space-based photobioreactors. This optical optimization provides the basis for the next step of this study: quantifying the electrical power needed to maintain such layers either in 1 g or 0 g conditions for potential space-based applications.

### Performance of the acoustic trapping at 0 g and 1 g

To determine the minimum electrical power required for stable trapping in microgravity, a flight-qualified millifluidic chip was boarded on the Novespace AirZero-G aircraft. Each parabola provides a 22 s 0 g (microgravity) phase, preceded and followed by 1.8 g pull-up and pull-out hypergravity phases. Before take-off, the chip was loaded with a *L. indica* suspension, and the transducer was activated at 2 MHz and 10 V to maintain the cells in the acoustic traps during the hyper-g phases. Once in the microgravity phase, the driving voltage (U_ac_) was set to the tested value. A frequency sweep from 1.8 to 2.2 MHz (450 ms cycle) was then applied, inducing axial oscillations of the layers if the ARF was sufficiently strong (Fig. 1). The axial position of the layer was extracted from video recordings (ImageJ) and converted into oscillation amplitudes using MATLAB, as illustrated in Fig. S2. Figure 7a shows a typical temporal oscillation of the z-position of a layer. The oscillation amplitude was defined as the mean peak-to-peak displacement (values below the pixel size of 12-µm were scored as zero in the subsequent analysis, see Methods). Supplementary Figure S5 illustrates the analysis protocol. Time-series were collected for each U_ac_ value, ranging from 0.3 to 10 V in microgravity, with more than twenty oscillations measured per voltage (11 s recordings). Repeated measurements enabled estimation of the mean amplitude with greater accuracy than the limit of detection. Identical experiments were also conducted in the laboratory, with voltages ranging from 1 V to 10 V.

**Figure 7:**
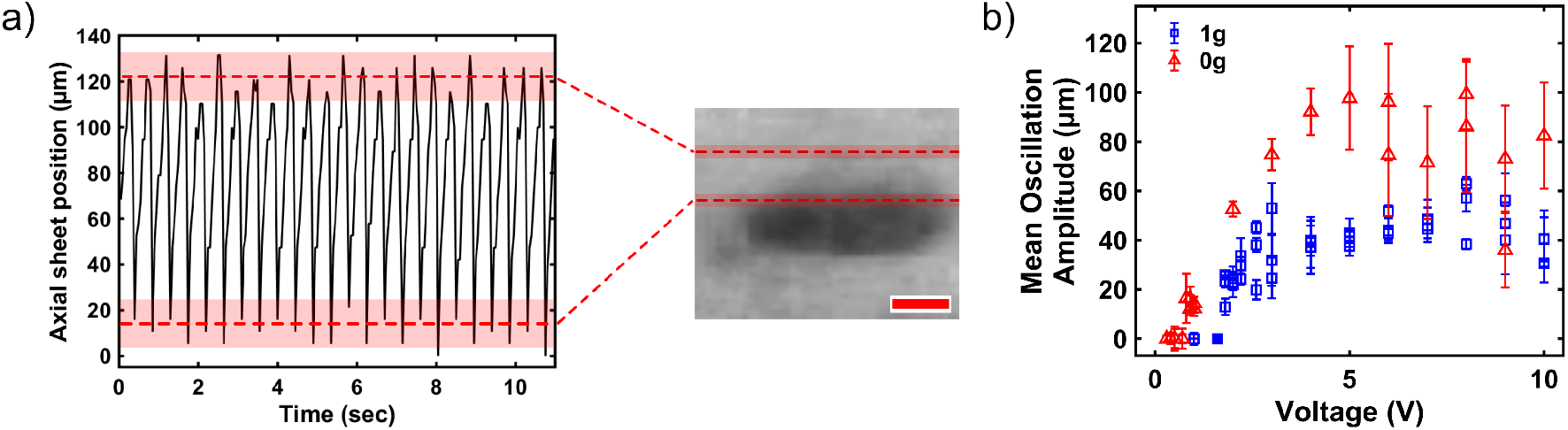
**a) Axial oscillations** of acoustically levitated *L. indica* PCC 8005 layers. Right: Cropped video frame (movie online) captured during a 0 g parabola, showing a cyanobacterial layer held at a pressure node. The red dashed lines show the extremal positions of the uppermost pixel of that layer during its up and down displacement. Left: Time evolution of the z-position of the uppermost pixel of that layer (*U*_*ac*_ = 5 V). The black curve shows the instantaneous position of the layer, while the red dashed lines mark the mean of the maxima and minima. The red shaded bands correspond to the standard deviation around those means. Scale bar is 100 µm. **b) Mean oscillation amplitude as a function of the voltage.** Blue squares represent the 1 g condition, while red triangles correspond to 0 g conditions. The points are the mean amplitude oscillation over more than 20 oscillations, while the error bars are the standard deviation around the mean.

Figure 7b shows the mean oscillation amplitude as a function of the applied voltage (Supplementary Fig. S3 presents the evolution of the oscillation amplitude as a function of the square of the voltage, which is proportional to the ARF). In microgravity, the first detectable oscillations appeared at 0.8 V. Their amplitude increased quasi-linearly up to 4 V, before saturating at a maximum of 100 µm for higher voltages. In laboratory conditions, oscillations began at 1.8 V. After a brief linear growth, the amplitude also saturated, but at a much smaller value of approximately 50 µm. For voltages below 1.8 V, gravity dominated over the ARF, causing sedimentation of the layers. The electrical power corresponding to each voltage was measured (Supplementary Fig. S4a, yielding minimum power consumption values of 0.42 mW in microgravity and 1.4 mW in laboratory conditions.) Although both values are very low, power consumption on Earth is three times higher than in microgravity. This low electrical power requirement is a major advantage for space-based photobioreactor applications.

However, a potential concern is that the low power consumption observed may simply reflect the small dimensions of the initial chip. Since scalability is a critical issue for future applications, we addressed this question experimentally by testing a larger PDMS chip with an internal height of *h*_1_ = 8 cm^24^, while employing the same piezoelectric transducer under 1 g laboratory conditions. This chip contained 8 cm^3^ of culture, i.e. 20 times the volume of the 0.4 cm^3^ chip used in microgravity, while maintaining the same cell density (OD_750_ = 0.2). Layers of levitating cyanobacteria were successfully obtained at a minimum driving voltage of 1.8 V, as shown in Fig. S7a. The corresponding electrical power (Fig. S7b) remained essentially unchanged compared to the small chip, despite the 20-fold increase in culture volume. This result suggests that the electrical power required for acoustic trapping scales only weakly with the chamber height.

Overall, these results demonstrate that acoustic trapping of cyanobacterial layers requires very little electrical power in microgravity, and that this requirement remains essentially constant when scaling up the size of the culture chamber. This opens avenues to design large-scale, energy-efficient photobiore-actors for space missions.

## Discussion

In this study, the potential of acoustic levitation to handle and organize living, spiral *L. indica* filaments in both 1 g and 0 g conditions was evaluated. More importantly, the feasibility of forming stable layers of cyanobacteria separated by clear gaps that allow light to penetrate deeply into the culture was explored.

Acoustic layering mitigates the bottleneck effect induced by the *compensation point*, which corresponds to the fluid depth where illumination is insufficient for photosynthesis to meet the microorganisms’ energy requirements. Each layer can be uniformly illuminated, ensuring that most cells receive sufficient light. In contrast, conventional photobioreactors rely on constant agitation, which still leaves inner regions deprived of light^49^. Notably, the electrical power required to trap cyanobacterial clusters decreases from 1.4 mW at 1 g to 0.42 mW in microgravity. A small piezoelectric transducer can therefore operate continuously in space without significantly burdening available power. By comparison, a 12 V motor at 15 rpm consumes ∼300 mW to stir 1 l of culture. Moreover, laboratory experiments confirmed that increasing the culture volume twentyfold (from 0.4 to 8 cm^3^) with the same transducer does not increase electrical power consumption, indicating weak scaling of power with reactor height. In addition, since sedimentation is absent in microgravity, layers can remain stable long after once the acoustic field is switched off, further reducing energy demand.

Light management remains a key factor for photobioreactors. Simulations show that, in the layered configuration, about 67% of incident light exits through the *x* = 0 and *x* = *L*_*x*_ boundaries, with only 33% absorbed by the layers, whereas bulk cultures absorb nearly all incident light within a few centimetres. Light loss could be mitigated with reflective walls, while angled beam illumination offers better coverage than isotropic lighting by reducing inter-layer shadowing. As the cyanobacterial compensation point^16^ is typically ∼2 W·m^−2^ and the simulated fluence rates are normalized, incident intensity in future experiments can be tuned so that absolute fluence exceeds this threshold throughout the layered culture to sustain photosynthesis.

Conventional agitation not only distributes light but also renews nutrients and prevents local O_2_ buildup. In simulated microgravity experiments with *L. indica*, Ellena *et al*. reported that the absence of convection produces a thick, stagnant boundary layer where photosynthetically produced O_2_ accumulates, slowing growth^50^. Within the acoustofluidic device, cells are immobilized in layers and can be gently perfused. For example, with the oxygen diffusion coefficient 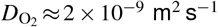 and a characteristic layer length *L* = 1 cm, a flow speed *v>D/L* ≈ 200 µm· s^−1^ yields a Péclet number 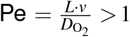, meaning convection outcompetes diffusion. In practice, perfusion enables O_2_ removal and nutrient renewal without mechanical stirring or filters, a key advantage for space missions where consumables are limited. Furthermore, although cyanobacteria tend to aggregate into dense layers, several parameters can be tuned to enhance nutrient access. For instance, introducing a frequency sweep can reposition layers and induce micro-stirring, improving nutrient transport and limiting local O_2_ buildup.

Refined modelling is required to predict ARF on helicoidal cyanobacteria. Existing theories mostly address spheres or simple elongated shapes^30,31,33,51^. Metallic nanorods can exhibit self-acoustophoresis in levitation planes^32,52^, but such effects occur due to the much lower density contrast compared to cyanobacteria. Furthermore, prior acoustofluidic studies in microgravity showed that frequency sweeps can displace layers but may induce Rayleigh–Bénard convection at 1 g due to transducer heating and this effect is suppressed in 0 g^40^. No convection-driven instabilities were observed in the present experiments, in either 1 g or 0 g.

Another uncertainty concerns the acoustic contrast factor of living cyanobacteria. Because these organisms can form gas vacuoles, it is unclear whether all trichomes localize at pressure nodes or antinodes, and this positioning may vary with physiological state. Under our experimental conditions, Fig. 5d shows that PCC8005 trichomes are trapped at pressure nodes. Longer-term studies will be required to probe state-dependent shifts. Nevertheless, whether cyanobacteria occupy nodes or antinodes does not affect the layered architecture, which consistently enhances light penetration.

Although acoustic energy density was not measured here, previous work using the same chip and transducer class (e.g., Jeger-Madiot *et al*.^43^, Fig. 18) established a clear relationship between input voltage and ARF magnitude, with ARF 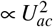. Thus, input voltage can serve as a practical proxy for trapping strength during frequency sweeps. While ARF peaks at resonance^18^, multiple power-consumption maxima were observed at 1.85, 2.00, and 2.15 MHz (Fig. S6), consistent with secondary ARF maxima in Jeger-Madiot *et al*.^43^. These maxima ensure that layers periodically experience sufficient ARF during sweeps. Since secondary-resonance ARF is weaker than the primary resonance, the sweep-based threshold is conservative, meaning the true voltage (power) threshold may be even lower.

Beyond growth, harvesting and filtering are also important for applications. Acoustic forces can accelerate sedimentation^24^ and have been used for biomass separation in H-shaped resonators under both terrestrial gravity and microgravity^34^. Because acoustic trapping concentrates and positions biomass without consumable membranes, it can reduce or eliminate the need for disposable filters, lowering launch mass critical for space missions and ensuring long-lasting, efficient recycling systems.

Scaling up to larger photobioreactors (e.g., 50 cm depth) will require careful power budgeting. Future studies will investigate how reactor dimensions affect photosynthetic efficiency, oxygen production, and nutrient availability with acoustic layering. A complementary long-term experiment (full growth curves under continuous levitation) is underway and will report physiological endpoints separately, while the present work establishes trapping feasibility, light penetration, and power cost.

Altogether, layered cultures reduce energy demand and mechanical complexity while lowering contamination risks associated with moving parts. Acoustic structuring thus emerges as a promising strategy for sustainable biomass production and CO_2_ recycling in long-duration space missions.

## Code and data availability

All relevant data and codes are available upon demand.

## Aknowledgement

The authors wish to thank ESA and the Carnot IPGG for funding the PhD thesis of B. Dupont. The participation in the parabolic flights campaign was financially supported by the CNES (French Aerospace Agency). The authors thank Corinne and Franck Chauvat (CEA Saclay) for their advises on cyanobacterial culture and insightful discussions. The authors thank G. Gauquelin-Koch (in charge of the Life Science topic at the CNES) and T. Bret-Dibat (in charge of the Matter Science topic at the CNES). The authors also thank T. Paris (Novespace) for his help in optimizing the experimental setup to make it fit the constraints of the Airbus Air Zero-G. The authors finally thank J.-M. Peyrin (Neurosciences Paris Seine, Sorbonne Université) and P. E. Lecoq (NPS and PMMH) for their help during the preparation of the setup and the flights.

## Author contribution

B.D., X.B-G., J-L.A. and M. A.: conception and fabrication of the setup. B.D., J-L.A. and M.A.: conceptualization and methodology. B.D., J.-L.A. and M.A.: samples preparation, data acquisition, analysis and simulations. B.D., S.B-V., J.-L.A. and M.A.:writing original draft, review and editing. S.B-V., J.L.A. and M.A.: supervised the project. All authors contributed to the article and approved the submitted version.

## Supplementary data

### Supplementary movies

Video associated with the figure 5. Formation of layers of cyanobacteria (*L. indica* PCC 8005) under stationary acoustic field. Scale bar = 0.5 mm.

Video associated with the figure 7. Side observation showing the axial oscillations of acoustically levitated *Limnospira indica* PCC 8005 layers during microgravity phase (0 g). Scale bar = 300 µm. Media: https://figshare.com/s/9dd41098f2ccb29c8522

### Cell viability after exposure to acoustic levitation

To validate that acoustic levitation does not harm *L. indica* PCC 8005 cells, growth before and after acoustic exposure was compared. For this purpose, a custom-designed stainless-steel resonant cavity (55 mm × 55 mm × 55 mm), equipped with watertight lateral poly(methyl methacrylate) (PMMA) windows and a threaded ring to mount a large ultrasonic transducer (Olympus™, model A395S), was used.

Each experiment lasted one week. Acoustic exposure was applied for 6 h at room temperature (shaded area in Fig. S1a,b). Following acoustic exposure, cells were transferred to Erlenmeyer flasks and incubated at 27 °C in a rotary shaker (140 rpm) under a 12 h light /12 h dark cycle, with a light intensity of approximately 6.64 W·m^−2^ (≈ 2500 lux). Optical density at 750 nm (OD_750_) was recorded at 0 h (beginning of levitation), 6 h (ending of levitation), 48 h, and 168 h.

Two experimental series were conducted:

1. First series (Fig. S1a): two replicates of levitated cultures and one non-levitated control that was placed in the stainless-steel resonant cavity and then transferred to an Erlenmeyer flask.
2. Second series (Fig. S1b): two replicates of non-levitated cultures placed in the stainless-steel resonant cavity and then transferred to Erlenmeyer flasks; three replicates maintained in Erlenmeyer flasks at room temperature for 6 h (not exposed to the cavity); and one levitated culture placed in the stainless-steel resonant cavity and then transferred to an Erlenmeyer flask.

Growth curves revealed no adverse effects from acoustic exposure: levitated cultures grew comparably to the controls.

In addition, microscopic images of the cyanobacteria were acquired (Zeiss microscope, 5× objective) to visually assess morphology. Fig. S1c shows representative trichomes before acoustic levitation; Fig. S1d shows trichomes after 1 h of acoustic levitation; and Fig. S1e shows trichomes from a control sample after 1 h of growth without acoustic levitation. No discernible morphological differences were observed between levitated and non-levitated cultures.

**Figure S1:**
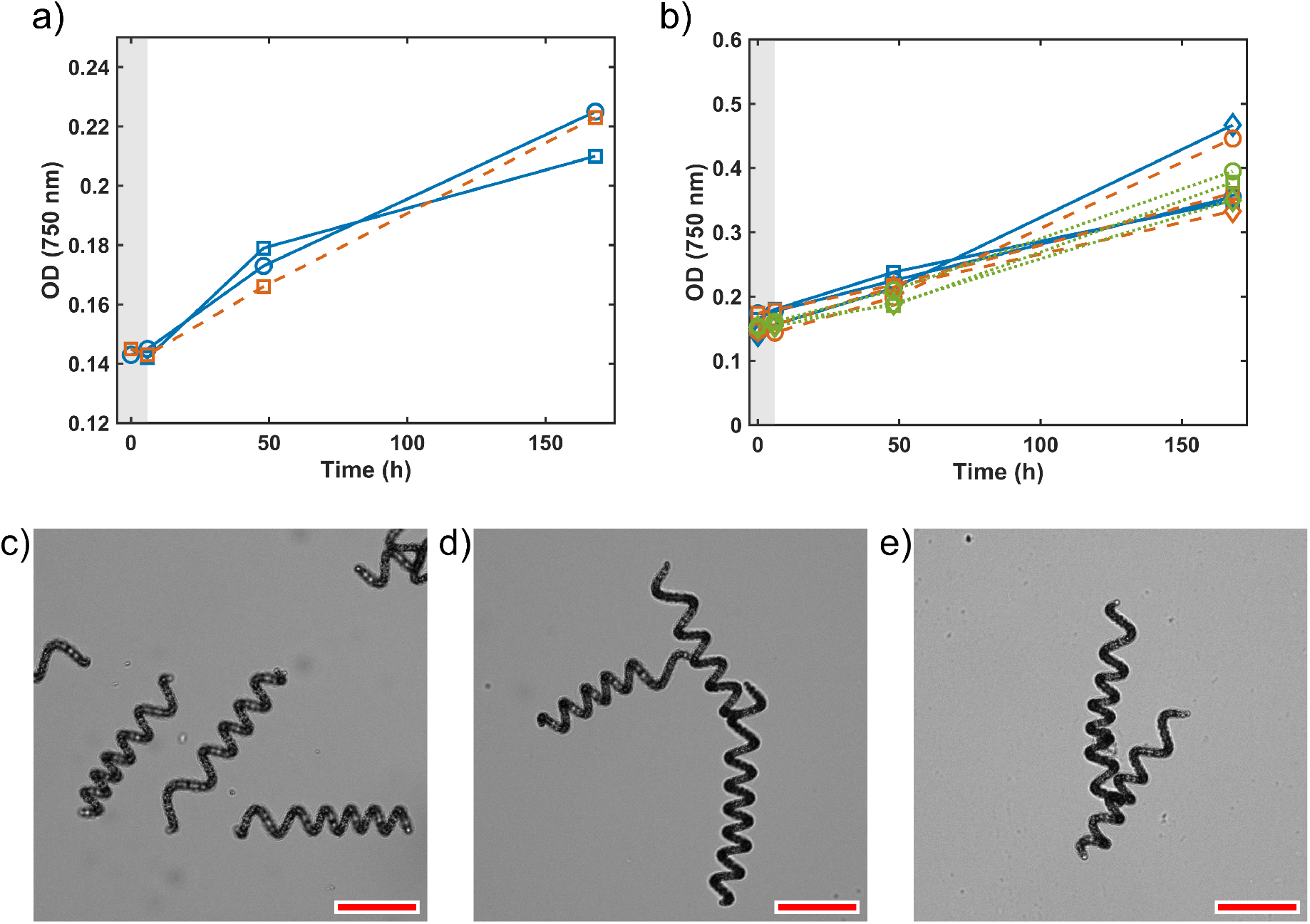
Monitoring the viability of *Limnospira indica* PCC 8005 under different culture conditions. Comparison between levitated and non levitated cyanobacteria. Blue curves correspond to levitated cell cultures, orange curves to non-levitated cultures, and green curves to cultures kept at room temperature and agitated at 140 rpm in a rotary shaker incubator for 6 hours in an Erlenmeyer instead of being in the stainless steel cavity. Each replicate is indicated by a distinct symbol: circle, square, or diamond. Acoustic levitation was applied between 0 and 6 hours (grey shaded area), after which cultures were monitored under standard growth conditions at 48 and 168 hours. **a)** First series of test. **b)** Second series of test. **c)** Representative image of trichomes before acoustic levitation. **d)** Representative image of trichomes after to 1h of acoustic levitation. **e)** Representative image of trichomes after to 1h in the levitation device but with the acoustic OFF.

**Figure S2:**
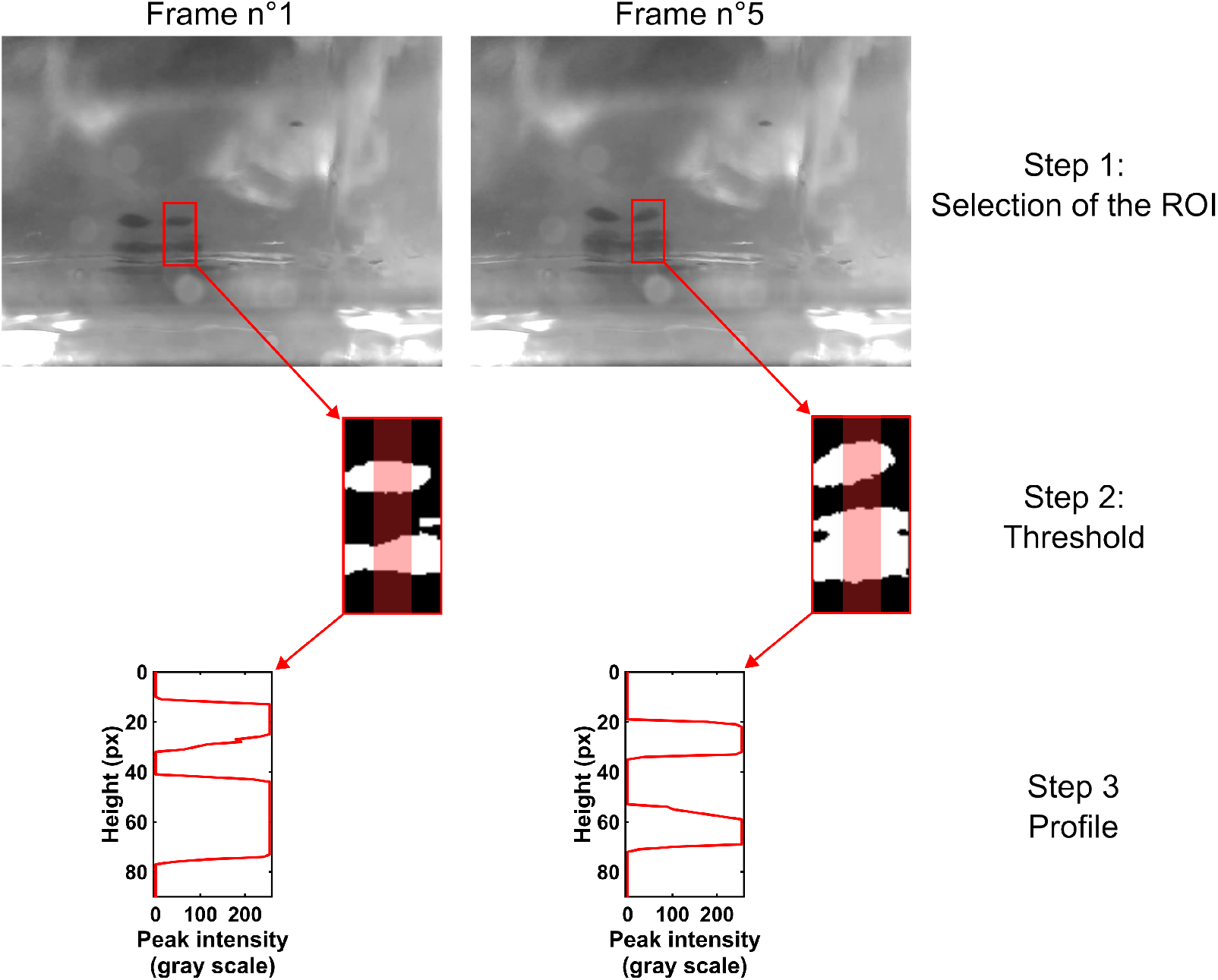
Processing steps to determine the mean amplitude of oscillation of a sheet under acoustic levitation. **a)** Images (8-bit grayscale) of the side view of the resonant cavity with the layers.**b)** Images cropped (45 px-wide × 90 px-high) and converted to binary by applying a threshold. **c)** Vertical profiles are extracted along the red line, with pixel intensities averaged across its width. The oscillation amplitude is then determined from the position of the first pixel, starting from the top of the profile, that has an intensity value of 255.

**Figure S3:**
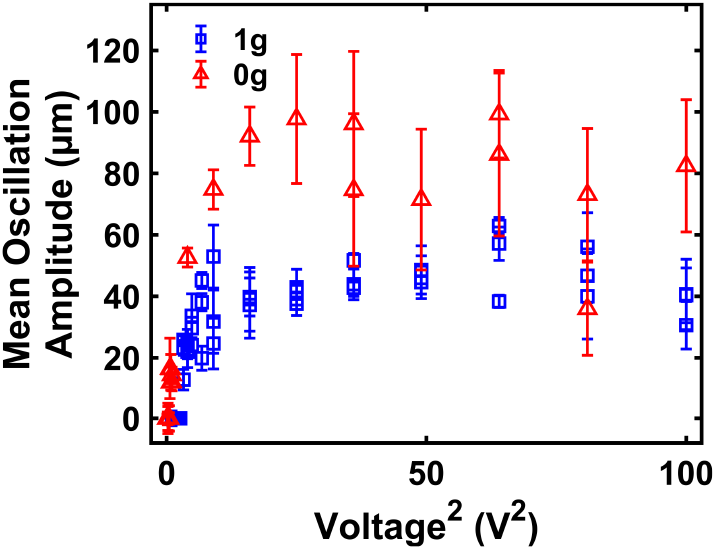
Mean fluctuations of *Limnospira indica* PCC 8005 sheet under acoustic levitation as a function of square of the voltage. The blue squares represent the mean oscillations of the sheet under 1 g conditions, whereas the red triangles indicate the mean oscillations of the sheet under 0 g conditions.

**Figure S4:**
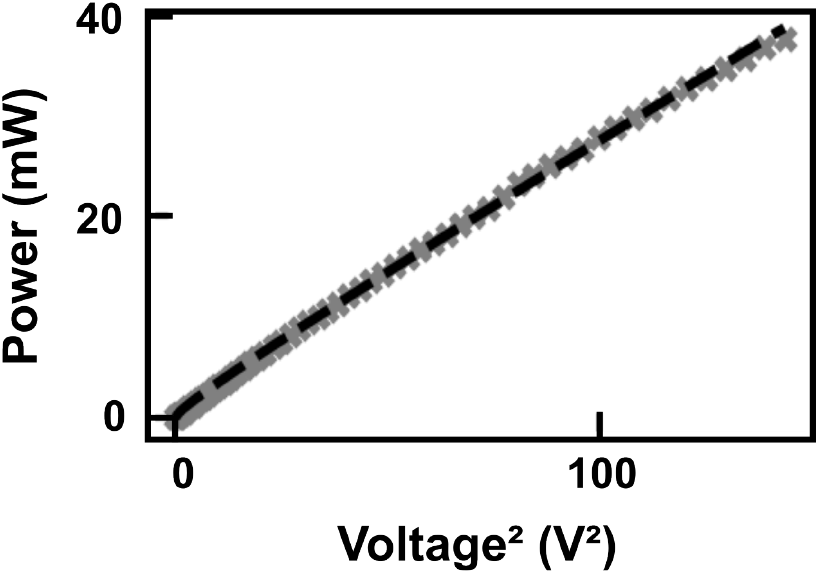
Correlation between the square of the applied voltage and the measured electric power. Gray crosses are experimental values while the black dashed line is the linear fit. The power is fitted by *P* = 0.24 · *V* ^2^ + 0.33 · *V*.

**Figure S5:**
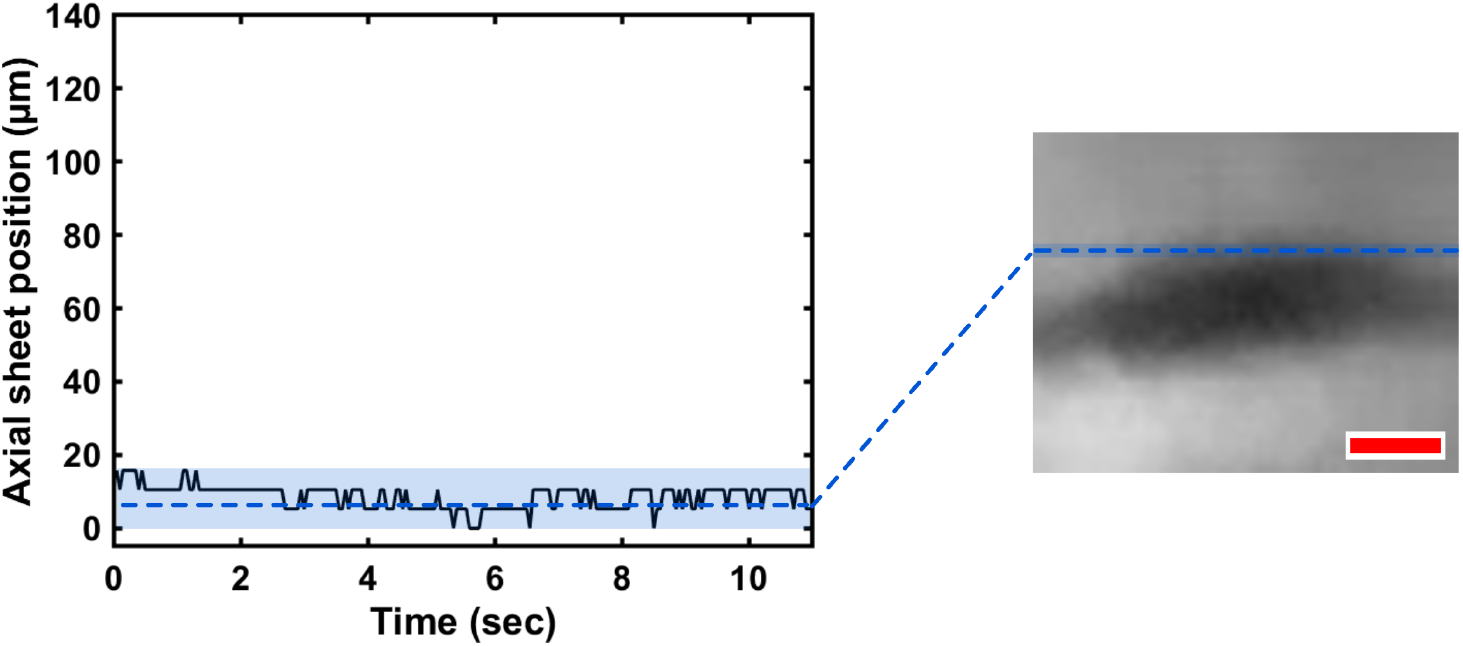
Example of a non-oscillating aggregate at 0.3 V under microgravity. The black curve shows the evolution of the vertical position of the aggregate over time, in microgravity (5 V). Blue dotted lines indicate the average value of the detected peaks of the respective maximum and minimum displacement amplitudes. The blue shaded region represents the standard deviation of the aggregate’s oscillations around the averaged values. Scale bar is 100 *µ*m.

**Figure S6:**
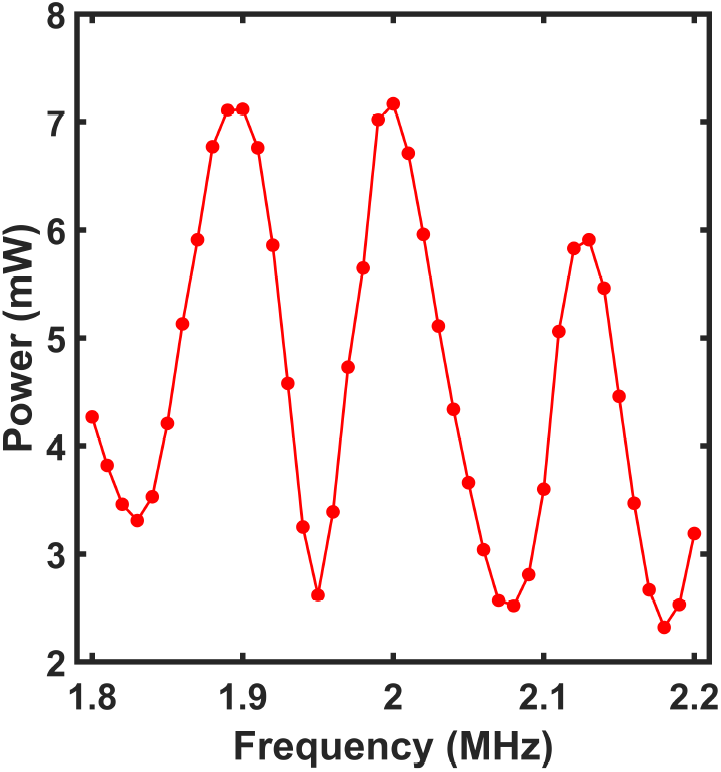
Electric power measurement as a function of the acoustic frequency with *L. indica* PCC 8005 suspension. The power is measured for a frequency ranging between 1.80 and 2.20 MHz (5 V). Measurements were carried out using a suspension of *L. indica* PCC 8005 with an optical density (OD_750_) of 0.2.

**Figure S7:**
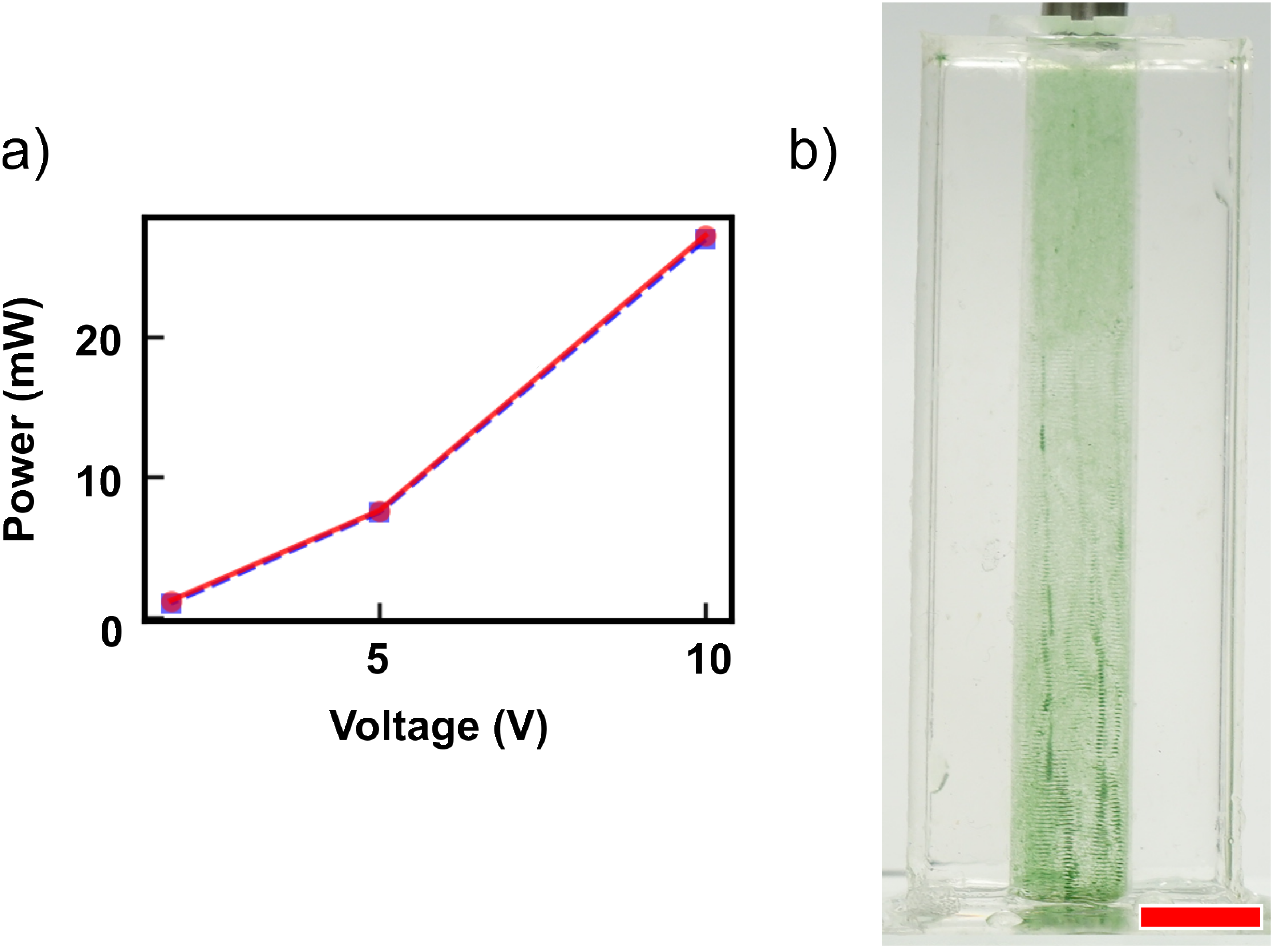
Scale-up of the acoustic levitation cavity. **a)** Electrical power consumed by the piezoelectric transducer for three voltages (1.8 V, 5 V and 10 V) in a 4 mm-high chip (red) and an 8 cm-high chip (blue). **b)** Acoustic trapping of *L. indica* PCC8005 in the 8 cm-high chip at 2.00 MHz and 1.8 V (minimum voltage to obtain levitation in the laboratory). Scale bar is 1 cm.

**Figure S8:**
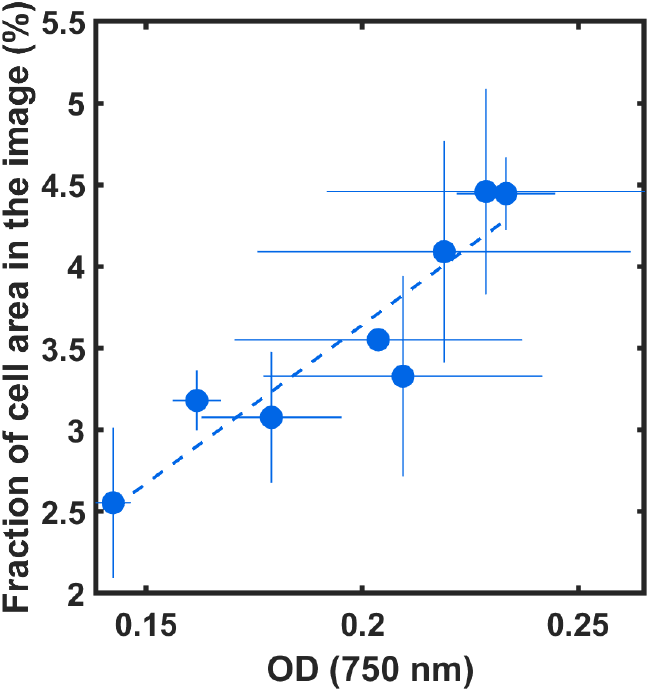
Linearity between optical density at 750 nm (OD_750_) and percentage of trichome’s area in photos. OD measurements were performed on three independent replicates, each corresponding to a different time point during cyanobacterial growth. Each OD value represents the mean of three measurements. At the same time images are taken with a Malassez counting chamber using a Zeiss microscope at 5x magnification. The fraction of area corresponds to the percentage of image pixels occupied by cyanobacteria. It is calculated as the ratio of cell-occupied pixels to the total number of pixels in the image. For independent replicate, the measure is done on three samples. This, each value of the fraction area is the mean of 6 measurements. Error bars indicate the standard deviation for both OD and area measurements. A linear regression (dashed line) is shown. The linear fit is depicted by the blue dashed line: *y* = 19.397 × *x* − 0.237

**Figure S9:**
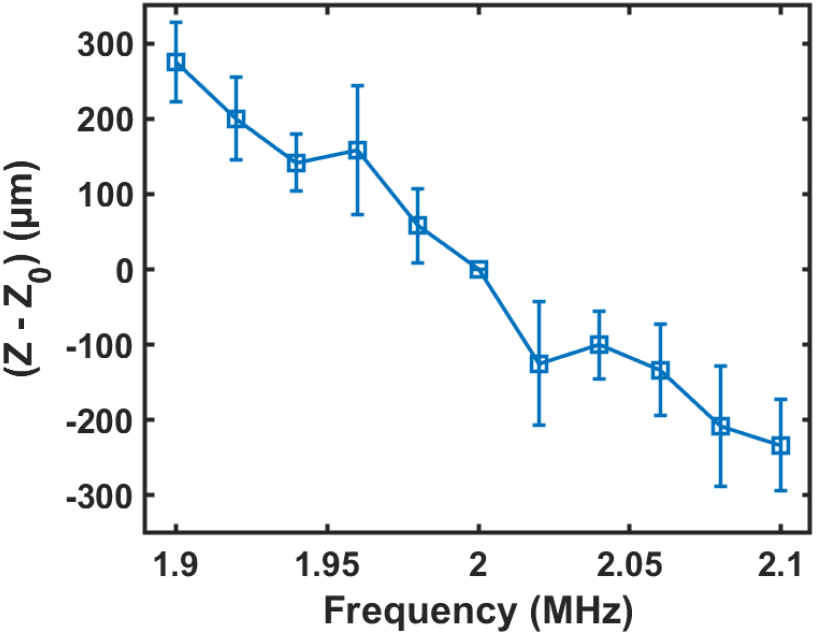
Axial displacement of levitated cyanobacterial layers as a function of acoustic frequency. Levitated *L. indica* PCC8005 culture (OD_750_ = 0.085) in the millifluidic chip (cavity height = 3 mm), recorded during a frequency sweep from 1.90 to 2.10 MHz at 10 V. Blue squares represent the mean axial (*z*) position of three levitated layers. Error bars indicate the standard deviation across three layers in two independent replicates. The axial position is normalized by the reference position *z*_0_ at 2.00 MHz to highlight the relative amplitude of displacement.

